# Activational and organizational effects of sex hormones on hippocampal inhibitory neurons

**DOI:** 10.1101/2024.01.20.576232

**Authors:** Alicia Hernández-Vivanco, Rut de la Vega-Ruiz, Alberto Montes-Mellado, Íñigo Azcoitia, Pablo Méndez

## Abstract

Peripheral and brain-produced sex hormones exert sex-specific regulation of hippocampal cognitive function. Estrogens produced by neuronal aromatase regulate inhibitory neurons (INs) and hippocampal-dependent memory in adult female mice, but not in males. How and when this sex effect is stablished and how peripheral and brain sources of estrogens interact in the control of hippocampal INs is currently unknown. Using ex-vivo electrophysiology, fiber photometry, molecular analysis, estrous cycle monitoring and neonatal hormonal manipulations, we unveil estrous cycle dependent and independent features of CA1 Parvalbumin (PV) INs and hippocampal inhibition in adult female mice. Before puberty, aromatase is expressed in PV INs and regulates synaptic inhibition in female but not in male mice. Neonatal testosterone administration altered prepubertal female mice hippocampal dependent memory, PV IN function and estrogenic regulation of adult female synaptic inhibition and PV INs perineuronal nets. Our results suggest that sex differences in brain-derived estrogen regulation of CA1 inhibition are established by organizational effects of neonatal gonadal hormones and highlight the role of INs as mediators of the sexual differentiation of the hippocampus.

**Highlights:**

- Estrous cycle dependent and independent features of CA1 PV INs and hippocampal inhibition
- Aromatase is expressed in male and female PV neurons before puberty.
- Neuroestrogens regulate prepubertal CA1 synaptic inhibition in females but not in males.
- Neonatal testosterone disrupts neuroestrogen effects on adult female hippocampus.
- Neonatal testosterone affects PV INs and hippocampal function before puberty.

## Introduction

In the adult brain, sex hormones regulate neuronal function and influence cognition through sex-specific actions in the hippocampus (Fleischer and Frick, 2023), a brain structure involved in learning, memory and spatial navigation. Sex-specific hormonal effects support basic neuronal mechanisms underlying cognitive function (Azcoitia et al., 2022; Yagi and Galea, 2019) and have been linked to the sex bias in the prevalence of neurodevelopment disorders, such as Autism Spectrum Disorders and Intellectual Disability (Bölte et al., 2023). Moreover, changes in sex hormone levels and reproductive function interact with aging in promoting cognitive deficits (Crestol et al., 2023; Lopez-Lee et al., 2024; Zárate et al., 2017). Despite the relevance to understand cognition in the healthy, aging or diseased brain, how and when sex effects are implemented in the hippocampus is not fully understood.

Estrogens regulate the function of hippocampal Gamma-Amino Butyric Acid (GABA)-releasing inhibitory neurons (INs) (Huang and Woolley, 2012; Murphy et al., 1998). INs dictate the temporal coordination of excitatory neuronal activity underlying hippocampal cognitive functions (Klausberger and Somogyi, 2008). Estrogens reduce inhibitory neurotransmission onto CA1 excitatory pyramidal neurons (Huang and Woolley, 2012; Tabatadze et al., 2015), a process in which local production by neuronal aromatase is critically involved. Neuron-derived estrogens (neuroestrogens) reduce the coverage of hippocampal CA1 parvalbumin (PV)-expressing INs by perineuronal nets (PNNs), extracellular proteoglycan structures that enwrap PV INs and regulate excitability and plasticity of this IN type (Hernández-Vivanco et al., 2022). Importantly, akin to previous results on excitatory synaptic function (Kretz et al., 2004; Wang et al., 2018), neuroestrogen regulation of synaptic inhibition and PV-INs PNNs is only detected in female mice and not in males (Hernández-Vivanco et al., 2022; Huang and Woolley, 2012). The mechanisms that give rise to sex differences in the regulation of hippocampal PV INs by neuroestrogens are currently unknown.

Sex differences in mammalian brain arise from the different sex chromosome complement (XX and XY) of male and female neurons and from the action of local and peripheral produced sex hormones (McCarthy et al., 2012). The organizational -activational theory of brain sexual differentiation (Arnold, 2009) posits that hormonal production by late embryonic and neonatal testis trigger sex-specific genetic programs (Gegenhuber and Tollkuhn, 2020) with enduring consequences in connectivity and physiology of neuronal networks (organizational effects). In addition, activational effects of sex hormones released by the gonads after puberty exert transient and reversible actions in a sex specific manner (Arnold, 2009). Using genetically modified mice to break the link between gonadal and genetic sex, we have previously shown that neuroestrogens reduce CA1 synaptic inhibition and PV INs PNNs coverage in gonadal female mice with XX or XY sex chromosome complement (Hernández-Vivanco et al., 2022). These results suggest that female-specific neuroestrogens actions on hippocampal inhibition and PV INs PNNs are independent of the genetic sex of the brain and raise the alternative possibility that sex effects are determined by adult (activational) or neonatal (organizational) actions of gonadal hormones.

Here we investigated the origin of sex differences in estrogenic regulation of CA1 synaptic inhibition and hippocampal PV INs using ex-vivo electrophysiology, fiber photometry, molecular analysis, and estrous cycling monitoring. We first tested whether estrous cycle-related activational effects of ovarian hormones regulate CA1 synaptic inhibition, PV INs activity, PNNs and aromatase expression. We then determined whether neuroestrogen regulates CA1 synaptic inhibition before functional maturation of the gonads and used neonatal hormonal manipulations to test organizational effects on CA1 synaptic inhibition. Our results show estrous cycle dependent and independent features of CA1 PV INs and unveil organizational effects of neonatal gonadal hormones on hippocampal inhibition.

## Results

### Estrous cycle regulation of CA1 synaptic inhibition and PV INs

In order to investigate the effects of ovarian hormones on synaptic inhibition and INs in the adult hippocampus, we used slice electrophysiology to determine spontaneous Inhibitory Post-Synaptic Currents (sIPSCs) in CA1 excitatory pyramidal neurons, fiber photometry to record PV IN activity in vivo and histological analysis to measure perineuronal net (PNN) coverage of PV INs in the CA1 area in female mice in different stages of the estrous cycle. We chose those parameters because sIPSCs frequency, PV INs activity and PNNs coverage are increased by pharmacological reduction of neuroestrogen synthesis with aromatase inhibitors in female mice (Hernández-Vivanco et al., 2022). Additionally, we determined the influence of the estrous cycle on the expression of aromatase in CA1 PV INs. At 10-12 weeks of age, adult female mice were assigned to diestrus and proestrus groups by performing vaginal cytologies (Fig. 1A).

**Figure 1.**
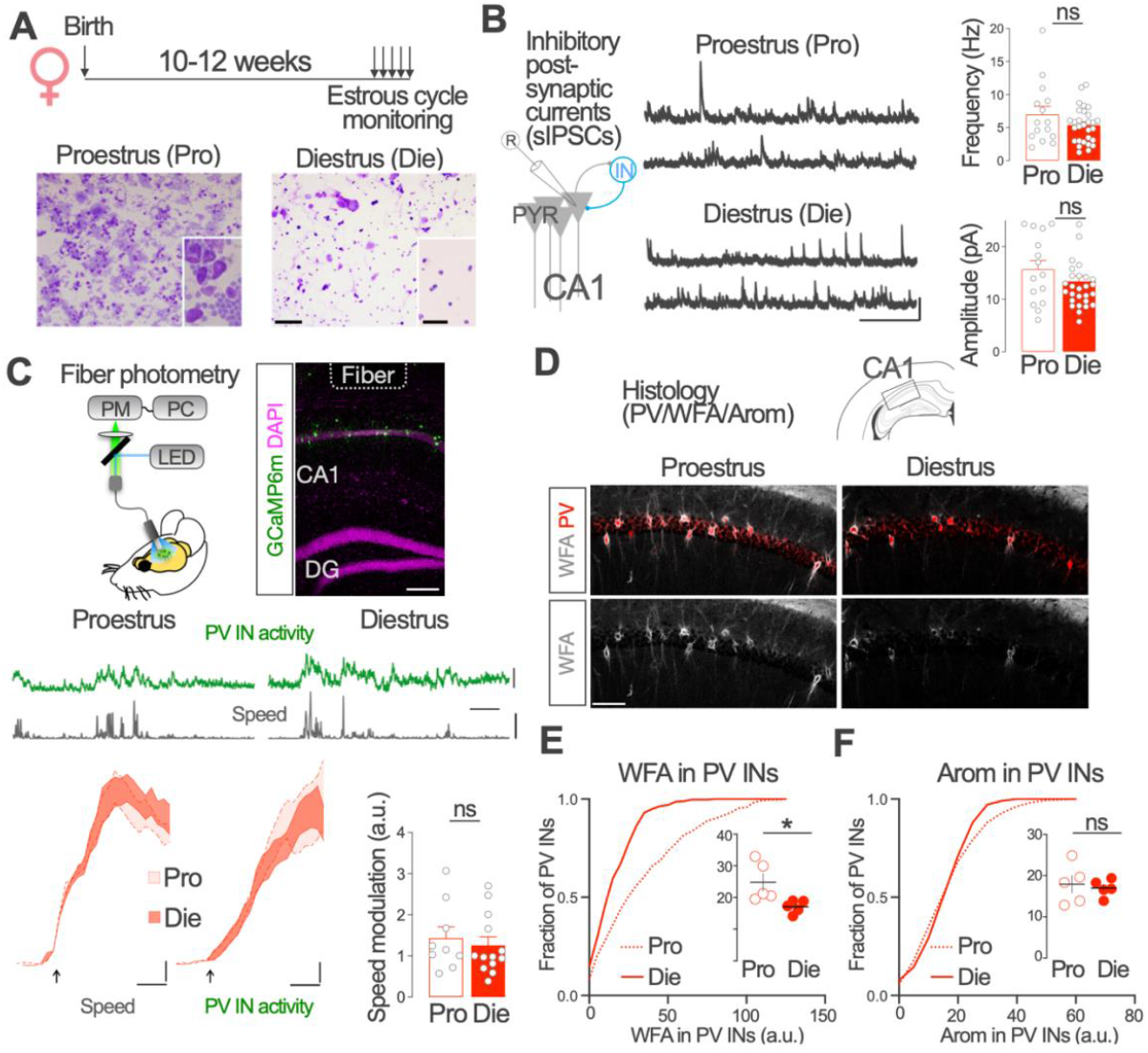
Estrous cycle-associated changes in CA1 synaptic inhibition and PV INs. **A**. Estrous cycle was monitored using vaginal cytologies in 10-12 weeks old female mice. Representative images (10 and 40 times magnification, inset) of cresyl violet-stained vaginal smears used to assign female mice to proestrus (left) and diestrus (right) stages of the estrous scycle. Scale bars: 100 μm (25 μm inset). **B**. In the morning of diestrus or proestrus, mice were processed for Spontaneous Inhibitory Post-Synaptic Currents (sIPSCs) recordings in acutely prepared brain slices. Representative recordings of sIPSCs in Proestrus (Pro) and diestrus (Die) female mice. Scale bar: 50 pA, 1 s. Graphs represent group data. Frequency, Two-tailed Mann Whitney test, U = 187, p = 0.46. Amplitude, Unpaired two tailed t-test, *t* (42) = 1.27, *p* = 0.21. *n* = 15, 29 neurons from 3-4 mice per group. **C**. Fiber photometry recordings were performed in adult female PV-Cre mice expressing GCaMP6m in dorsal CA1 PV INs using a chronically-implanted optic fiber while mice freely explore a familiar open field (upper panels). Scale bar: 200 μm. Mice speed (grey) and PV IN activity (GcaMP6m fluorescence, green) levels were simultaneously monitored in the morning of diestrus or proestrus stages of the estrous cycle (middle panels). Scale bars: 5 z-score, 5 cm/s, 30 s. Lower left plots show speed and z-scored PV IN activity (mean +/-SEM) aligned to locomotion onset (arrows) in proestrus and diestrus. Scale bars: 0.5 cm/s, 1 z-score, 0.5 s. Lower right graph shows no differences in the positive relationship between PV IN activity and mice speed in diestrus and proestrus. Two-tailed Mann Whitney test, U = 45, p = 0.39. *n* = 9, 13 recordings from 5 mice. **D**. Upper panels: representative image of simultaneous parvalbumin (PV, red) immunohistochemical detection and WFA staining of PNNs (grey) in dorsal hippocampus CA1 region of an adult female mice in proestrus (left) and diestrus (right). Single channel image of WFA staining is represented in grey in the lower part of the panel. Scale bar: 100 μm. Lower panel left: group data of WFA (left) and aromatase (right) staining intensities in PV-INs. Cumulative frequency distribution of individual values per PV IN and mean values per mice are shown. WFA: unpaired two tailed t-test, *t* (8) = 2.67, *p* = 0.03. Aromatase: unpaired two tailed t-test, *t* (8) = 0.39, *p* = 0.7. *n* = 5 mice per group. Graphs represent mean +/-SEM (columns and bars) and individual values (recorded neurons, grey circles in B and mice; circles in E). Graphs in E represent cumulative distribution of WFA and aromatase staining in individual PV INs.* p < 0.05; ns p > 0.05.

We recorded sIPSCs on visually identified CA1 pyramidal neurons with intact network activity in acutely prepared brain slices from female mice processed in proestrus or diestrus. In contrast to the previously observed regulation by neuroestrogen (Hernández-Vivanco et al., 2022), the frequency and amplitude of sIPSCs did not show apparent differences between proestrus and diestrus female mice (Fig. 1B).

We then used fiber photometry to record the activity of dorsal CA1 PV INs expressing the calcium sensor GCaMP6m in freely-moving adult female mice exploring a familiar enclosure. We simultaneously tracked mice speed in the enclosure and recorded calcium dependent GCaMP6m fluorescence (Fig. 1C, upper panels). We used the latter as a surrogate of PV IN population activity.

In agreement with previous reports (Arriaga and Han, 2019; Dudok et al., 2021; Hainmueller et al., 2024), we observed a strong coupling between PV IN activity and mice locomotion. Locomotion-associated changes in PV IN activity were evident in recordings obtained during both diestrus and proestrus stages (Fig. 1C, middle panels). By plotting the relationship between z-scored GCaMP6 fluorescence intensity and mice speed (Fig. 1C, lower panels), we observed no differences in locomotion-regulated PV IN activity between proestrus and diestrus.

Lastly, we used histological sections from diestrus or proestrus female mice to determine PNN coverage of CA1 PV INs (Wisteria floribunda agglutinin staining, see Methods) and aromatase expression in PV INs with specific antibodies. WFA intensity around PV INs in proestrus was higher compared with diestrus (Fig. 1D, E). In contrast, diestrus and proestrus female mice showed similar levels of aromatase expression in PV INs (Fig. 1F).

These results show that PV INs PNN coverage fluctuates across the estrous cycle, increasing during proestrus. Estrous cycle does not apparently modify synaptic inhibition in CA1pyramidal neurons, PV IN activity or aromatase expression in female mice PV INs. These data unveil estrous cycle-dependent and independent features of CA1 PV INs and hippocampal inhibition. Together with the previously observed effects of neuroestrogen (Hernández-Vivanco et al., 2022), these results suggest that ovarian hormones and neuroestrogen exert different and independent activational effects on CA1 synaptic inhibition and PV INs.

### Aromatase expression and neuroestrogen production in PV INs before puberty

The previous results suggest that neuroestrogen affects hippocampal INs independently of the function of the adult ovaries. To further test this idea, we investigated neuroestrogen production by hippocampal PV INs before puberty, i.e., before the start of adult gonadal hormone production in male and female mice. We used immunohistochemistry to detect aromatase protein in CA1 PV INs at postnatal day (PND) 21, before puberty onset in mice (Fig. 1A).

Aromatase expression was observed in CA1 region of PND21 male and female mice (Fig. 2B). Aromatase immunoreactive cells were found in different layers, mainly in the pyramidal and oriens strata (Fig. 2B). Simultaneous localization of aromatase and PV in CA1 area of male and female mice showed aromatase expression in this IN type in male and female mice (Fig. 2B). Aromatase expression levels in CA1 PV INs of PND 21 male and female mice did not show significant differences (Fig. 2C). WFA staining indicated that aromatase-expressing PV INs were surrounded by PNNs in male and female CA1 region at PND 21 (Fig. 2C).

**Figure 2.**
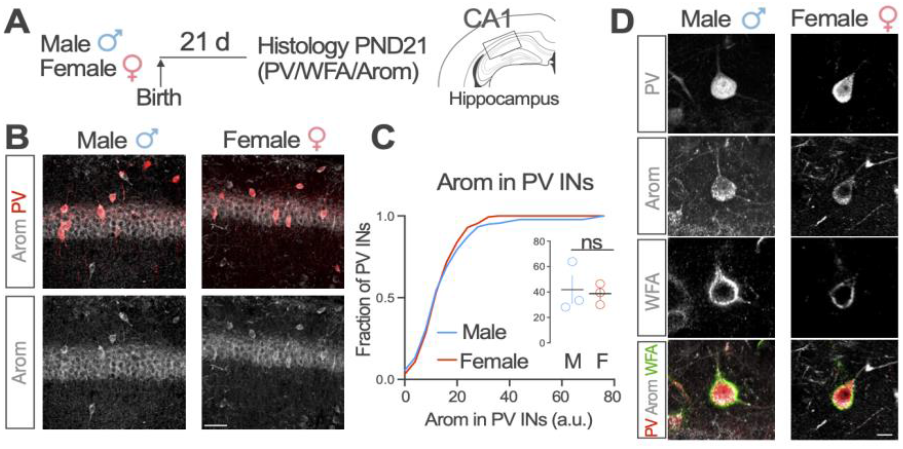
Aromatase expression in CA1 PV INs in prepubertal mice. **A**. Male and female mice were processed for immunohistochemistry at post-natal day 21. **B**. Representative immunofluorescence confocal microscopy images of aromatase (grey) and parvalbumin (red, upper panels) expression in the CA1 region of male (left) and female (right) hippocampus at 21 days of age. Single channel image of aromatase staining is represented in grey scale in the lower part of the panel. Scale bar: 50 μm. **C**. Quantification of aromatase expression level in M and F hippocampus. Group data of aromatase staining intensities in PV-INs. Cumulative frequency distribution of individual values per PV IN and mean values per mice are shown. Two-tailed Mann Whitney test, U = 4, *p* > 0.99. *n* = 3 mice per group. **D**. Representative immunofluorescence confocal microscopy images of simultaneous detection of parvalbumin (PV), aromatase (Arom) and and WFA staining of PNNs (WFA) in the CA1 region of male and female hippocampus at 21 days of age. Graphs represent mean +/-SEM and individual values. ns p > 0.05.

These results show that aromatase is expressed in PV INs before the start of sex hormone production by adult gonads in male and female mice and suggest prepubertal synthesis of neuroestrogen by hippocampal PV INs covered with PNNs.

### Estrogen regulation of CA1 synaptic inhibition before puberty

The presence of aromatase in PV IN at PND 21 suggest a functional impact of neuroestrogen on synaptic inhibition onto CA1 excitatory pyramidal neurons in prepubertal mice. To test this idea, we treated male and female mice with aromatase blocker Letrozole between PND21 and PND25 (0.5 mg/kg, one daily intraperitoneal injection during 5 days, Arom Block, Fig. 3A). Letrozole crosses the blood-brain barrier (Zhou et al., 2010) and has been previously shown to increase synaptic inhibition in adult female mice hippocampus (Hernández-Vivanco et al., 2022). On PND25, at the end of the treatment period, we performed patch-clamp recordings of sIPSCs from CA1 pyramidal neurons in acutely-prepared brain slices.

**Figure 3.**
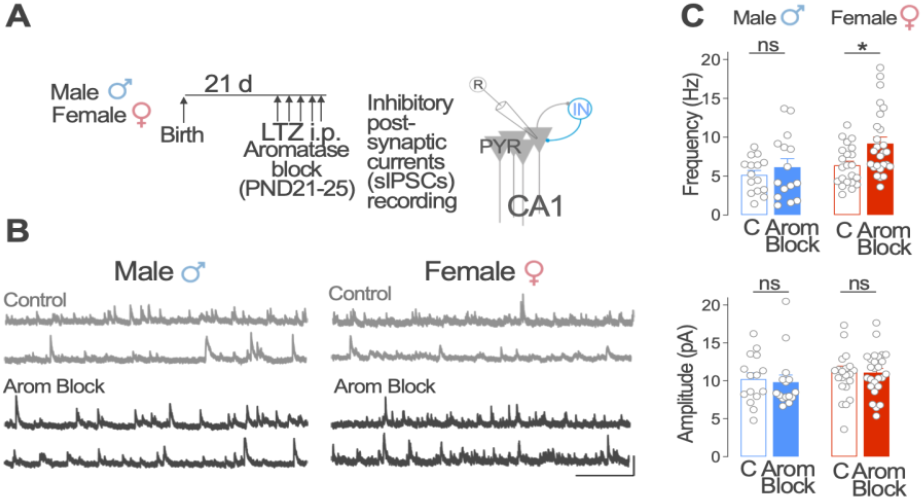
Aromatase regulation of CA1 synaptic inhibition in prepubertal male and female mice. **A**. Post-Natal Day (PND) 21 male and female mice received daily intraperitoneal (i.p.) injections of the aromatase blocker letrozole (LTZ) or vehicle (C) for 5 days. Spontaneous Inhibitory Post-Synaptic Currents (sIPSCs) were recorded from CA1 pyramidal (PYR) neurons in acutely prepared slices at PND25. **B**. Representative sIPSCs recordings from control (C, grey) and letrozole (Arom Block, black) treated male (left) and female (right) mice. Scale bar: 50 pA, 1 s. **C**. Group data from sIPSCs recordings. Frequency, two-way ANOVA, C/Arom Block *F* (1, 74) = 5.48, *p* = 0.02, Male/Female *F* (1, 74) = 6.9, *p* = 0.01. Bonferroni comparison tests, Male C vs Arom Block *p* = 0.87; Female C vs Arom Block *p* = 0.01. Amplitude, two-way ANOVA, C/Arom Block *F* (1, 74) = 0.09, *p* = 0.77, Male/Female *F* (1, 74) = 1.97, *p* = 0.16. Bonferroni comparison tests, Male C vs Arom Block *p* > 0.99; Female C vs Arom Block *p* > 0.99. Males, *n* = 15 neurons from 3 mice per group; females, *n* = 22 (Control), 27 (Arom Block) neurons from 3 mice per group. Graphs represent mean +/-SEM (columns and bars) and individual values (recorded neurons, grey circles). * p < 0.05; ns p > 0.05.

In female mice, aromatase blockade increased sIPSCs frequency in CA1 pyramidal neurons compared with vehicle treated mice (Fig. 3B,C). We observed no significant change in the amplitude of sIPSC in female mice (Fig. 3B,C). In contrast, in male mice, aromatase blockade did not produce apparent changes in sIPSC frequency and amplitude compared with vehicle treated male mice (Fig. 3B, C).

These results show that aromatase inhibition before puberty increases synaptic inhibition onto CA1 excitatory pyramidal neurons in female mice, but not in male mice.

### Neonatal testosterone impact on neuroestrogen regulation of CA1 synaptic inhibition and PV INs PNNs

The observed female-specific effects of neuroestrogen in prepubertal mice suggest that sex-specific neuroestrogen regulation of synaptic inhibition originates from early-life organizational effects of neonatal hormones. To test this idea, we treated neonatal female mice pups with testosterone propionate (100 μg in 50 μl of sesame oil, one daily injection on PND 1, 8 and 15) to mimic the male-specific perinatal testosterone surge. This treatment has been previously shown to masculinize behavior dependent on aromatase expressing neurons in different regions of the female brain (Wu et al., 2009). Ten weeks after testosterone or vehicle neonatal treatment, we tested neuroestrogen regulation of synaptic inhibition and PV INs PNNs by treating adult mice at 12 weeks of age with the aromatase specific inhibitor letrozole (0.5 mg/kg, one daily intraperitoneal injection during 5 days, Arom Block, Fig. 4A). We determined sIPSCs in CA1 pyramidal neurons and PV INs PNNs coverage using electrophysiological recordings and immunohistochemistry, respectively.

**Figure 4.**
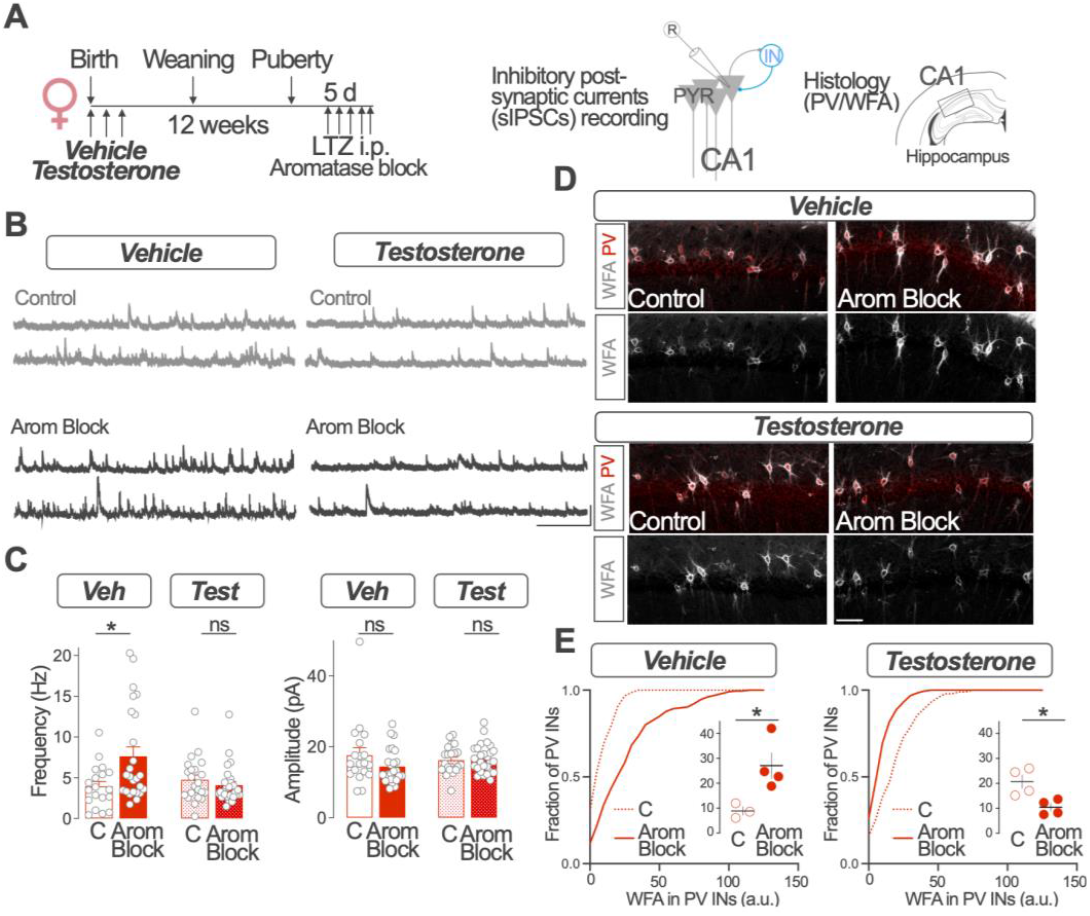
Neonatal testosterone effects on the regulation of hippocampal inhibition. Female mice pups received subcutaneous testosterone propionate (100 μg) or vehicle (sesame oil) on post-natal days 1, 7 and 15, weaned and raised to young adulthood (12 weeks). Adult mice were then treated with the aromatase blocker letrozole (LTZ) or vehicle (C) for 5 days and processed for Spontaneous Inhibitory Post-Synaptic Currents (sIPSCs) recordings or histological analysis. **B**. Representative sIPSCs recordings from control (C, grey) and letrozole (Arom Block, black) treated female mice with neonatal vehicle (left) and testosterone (right) treatment. Scale bar: 50 pA, 1 s. **C**. Group data from sIPSCs recordings. Frequency, two-way ANOVA, C/Arom Block *F* (1, 90) = 3.97, *p* = 0.05, Veh/Test *F* (1, 90) = 3.2, *p* = 0.07. Bonferroni comparison tests, Veh C vs Arom Block *p* = 0.002; Test C vs Arom Block *p* = 0.78. Amplitude, two-way ANOVA, C/Arom Block *F* (1, 90) = 1.82, *p* = 0.18, Veh/Test *F* (1, 90) = 0.02, *p* = 0.89. Bonferroni comparison tests, Veh C vs Arom Block *p* = 0.14; Test C vs Arom Block *p* = 0.99. Vehicle, control, *n* = 19 neurons, 3 mice; Arom Block *n* = 25 neurons, 4 mice; Testosterone, control, *n* = 22 neurons, 4 mice; Arom Block *n* = 28 neurons, 4 mice. **D**. Representative image of simultaneous parvalbumin (PV, red) immunohistochemical detection and WFA staining (grey) in the CA1 region of neonatal vehicle (upper panels) or testosterone (lower panels) treated female mice which received daily intraperitoneal (i.p.) injections of the aromatase blocker letrozole (LTZ, Arom Block, right panels) or vehicle (Control, C, left panels). Single channel image of WFA staining is represented in grey scale in the lower part of the panels. Scale bar: 100 μm. **E**. Group data of WFA staining intensities in PV-INs for neonatal vehicle (left) or testosterone (right) treated female mice. Cumulative frequency distribution of individual values per PV IN and mean values per mice are shown in each case. Vehicle, C vs Arom Block, unpaired two tailed t-test, *t* (5) = 2.91, *p* = 0.03. Testosterone, C vs Arom Block, unpaired two tailed t-test, *t* (6) = 3.32, *p* = 0.02. Vehicle, control, *n* = 3 mice; Arom Block *n* = 4 mice; Testosterone, control, *n* = 4 mice; Arom Block *n* = 4 mice. Graphs represent mean +/-SEM (columns and bars) and individual values (recorded neurons, grey circles in C mice; circles in E). Graphs in E represent cumulative distribution of WFA staining for individual PV INs.* p < 0.05; ns p > 0.05.

In line with previous results in adult mice (Hernández-Vivanco et al., 2022) and prepubertal mice (Fig. 3C), aromatase blockade increased sIPSC frequency in neonatal vehicle treated female mice. While neonatal testosterone treatment did not produce significant sIPSCs frequency changes when compared with vehicle treated mice, it completely prevented the effect of aromatase blocker letrozole on sIPSC frequency (Fig. 4B,C). We observed no significant differences in the amplitude of sIPSCs between the experimental groups (Fig. 4B,C).

Cumulative frequency distribution analysis of WFA staining intensities showed that aromatase blockade increases the intensity of WFA staining surrounding CA1 PV INs in neonatal vehicle treated mice (Fig. 4D,E1, left panel). In contrast, aromatase inhibition reduced WFA staining in PV INs of neonatal testosterone-treated mice (Fig. 4 D,E, right panel).

These results show that neonatal testosterone treatment in female mice prevents neuroestrogen effects on synaptic inhibition in CA1 pyramidal neurons and disrupts the regulation of CA1 PV INs PNNs. These results strongly suggest organizational effects of neonatal hormones in neuroestrogen regulation of CA1 synaptic inhibition and CA1 PV INs.

### Neonatal testosterone effects on prepubertal PV INs and hippocampal function

We next investigated whether neonatal gonadal hormones impact hippocampal function and PV INs before puberty. We studied the effect of neonatal testosterone treatment on behavior of PND21 female mice during the training and recall in contextual fear conditioning (CFC), a hippocampal-dependent associative memory task. We additionally determined the expression of the neuronal activity marker c-Fos and PNN coverage in PV INs by immunohistochemical analysis of mice processed 90 min after fear memory recall (Fig. 5A).

**Figure 5.**
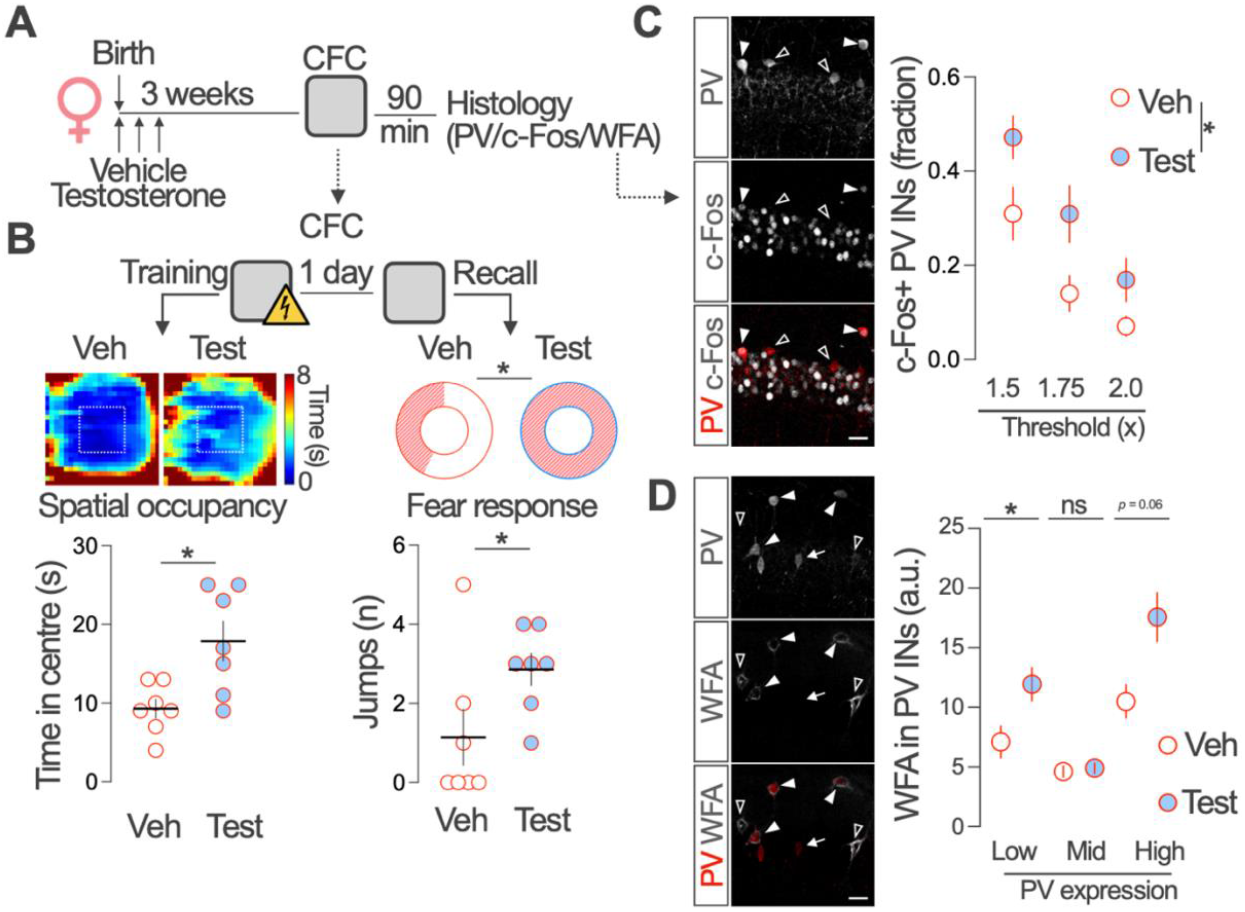
Neonatal testosterone regulates PV INs and hippocampal function in prepubertal female mice. **A**. Female mice pups received testosterone propionate (100 μg) or vehicle (sesame oil) on post-natal days 1, 7 and 15. On PND20, mice were trained in the Contextual Fear Conditioning (CFC) and, on PND21, tested for fear memory recall. Mice were processed for histological analysis of PV, c-Fos and WFA staining 90 min after the recall test. **B**. Left, group spatial occupancy maps during the exploration in CFC training session. Graphs represent group data for the time spent in the center area. Unpaired two tailed t-test, *t* (12) = 3.08, *p* = 0.009. *n* = 7 mice per group. Right, proportion of mice (donut plots) and number of climbs or jumps per animal (lower graph) during fear memory recall. STAT **C**. Representative image of simultaneous parvalbumin (PV, grey in upper panel, red in lower panel) and c-Fos (grey, middle and lower panel) immunohistochemical detection in the CA1 region of a testosterone treated female mice 90 min after fear memory recall. Graph compares the fraction of PV INs expressing cFos in vehicle and testosterone treated female mice using a dynamic threshold (1.5, 1.75, and 2 times background levels) analysis. Two-way ANOVA, Veh/Test *F* (1, 8) = 5.33, *p* = 0.04. *n* = 5 mice per group. **D**. Representative image of WFA (grey in middle and lower panel) staining around low (empty triangles), middle (arrows) and high (solid triangles) PV (grey in upper panel, red in lower panel) expressing IN in the CA1 region of a testosterone treated female mice. Kruskal-Wallis test, H = 138.9, *p* < 0.0001, Dunn’s multiple comparisons tests, Low, *p* < 0.0001; Mid, *p* = 0.92; High, *p* = 0.058; vehicle *n* = 55, 329, 64 and testosterone *n* = 73, 393, 52, for low, middle and high PV expression, respectively. Graphs represent mean +/-SEM and individual values (mice in B and C, PV neurons in D). * p < 0.05; ns p > 0.05.

During CFC training, mice freely explore the conditioning cage during 3 min before the delivery of electric shocks. Tracking mice speed and position during exploration revealed that, although vehicle and testosterone treated female mice explore the cage at similar speed (Veh, 2.52 ± 0.36 cm/s, testosterone 2.57 ± 0.49 cm/s, Mean ± SEM Two-tailed Mann Whitney test, U = 24, *p* > 0.99. *n* = 7 mice per group), female mice treated with testosterone spent significantly more time occupying the central area of the cage (Fig. 5B). Locomotory reaction to shocks did not differ in both groups of mice, suggesting no major differences in sensing the aversive stimulus (Veh, 16.0 cm/s, testosterone 17.4 cm/s, Two-tailed Mann Whitney test, U = 19, *p* > 0.54. *n* = 7 mice per group).

During fear memory recall, 24 h after training, we evaluated passive (freezing, immobility) and active (jumps and climbs) fear responses during the 5 min re-exposure to the conditioning context (no shocks). The total time spent in freezing behavior and immobility did not differ between groups (Freezing: Veh, 2.3 ± 1.3 s/min, testosterone 8.1 ± 6.7 s/min, Two-tailed Mann Whitney test, U = 22, *p* = 0.80. Immobility: Veh, 44.5 ± 1.3 s/min, testosterone 42.8 ± 2.5 s/min, Two-tailed Mann Whitney test, U = 19, *p* = 0.54, *n* = 7 mice per group). In contrast, the proportion of mice displaying active responses and number of climbs or jumps per animal were increased in the testosterone treated female mice group (Fig. 5B).

We used simultaneous PV and neuronal activity marker c-Fos immunohistochemistry of female mice processed after the recall session in combination with a multi-threshold analysis (see Methods) to asses PV IN activity during the recall session. This analysis revealed a higher fraction of c-Fos-expressing CA1 PV INs in testosterone treated female mice compared with vehicle treated female mice after fear memory recall (Fig. 5C). Although we observed similar density of CA1 PV+ and WFA+ INs in vehicle and testosterone treated female mice, testosterone treatment increased the intensity of WFA staining in PV INs expressing low but not middle levels of PV and showed a strong tendence to increase WFA staining intensity in cell expressing high levels of PV (Fig 5D).

These results suggest that neonatal testosterone treatment impacts hippocampal-dependent memory, increases the activity of PV INs during memory recall, and alters PV INs PNN coverage. Testosterone may in this way impact PV INs function and PNNs during the juvenile period and participate in the maturation and sexual differentiation of hippocampal networks.

## Discussion

Excitatory neurons are targets for activational actions of sex hormones in the female hippocampus (Taxier et al., 2020). Our results show estrous cycle related changes, as well as estrous cycle independent aspects of CA1 inhibition and PV IN activity. In particular, females in proestrus, a stage of the estrous cycle associated with a rise in circulating estradiol concentration, show increased PNN neuronal coverage in the dorsal CA1 hippocampus. In contrast, synaptic inhibition onto CA1 excitatory neurons, the main output target of PV INs, and locomotion related PV IN activity remain unaltered. In CA1, PNNs mostly surround a prominent type of PV INs, PV-expressing basket cells (Yamada and Jinno, 2015). Since PNNs regulate physiology and plasticity of PV INs (Fawcett et al., 2019), our results suggest that cyclic ovarian production of sex hormones may affect dorsal hippocampal function through the regulation of this PNNs in PV-expressing basket cells. The well documented involvement of hippocampal PV INs PNNs in learning and memory processes (Favuzzi et al., 2017; Ramsaran et al., 2023) suggest that PNNs modulation across the estrous cycle may be related with the effects of peripheral hormones on memory (Taxier et al., 2020) and on differential hippocampal engagement in spatial tasks (Korol et al., 2004). Additionally, fluctuation of PNNs across the estrous cycle may be responsible for the behavioral response to non-cognitive aspects such as stress or aging (Laham et al., 2022).

We have previously reported that, in adult female mice, reduced neuroestrogen levels increase PV INs PNN coverage (Hernández-Vivanco et al., 2022, see also Fig. 4E). In contrast, here we observed that during proestrus, a stage associated to high level of plasma and hippocampal estrogens (Kato et al., 2013), PNN coverage of PV INs is increased. The differential effect on CA1 PV INs PNNs suggests that neuron-derived estrogen and cycling gonadal-derived ovarian hormones regulate CA1 PV INs PNNs through different mechanisms. Estrous cycle regulation of PNNs may involve the actions of other ovarian derived hormones such as progesterone (Laham et al., 2022). Moreover, estrous cycle may affect the molecular composition of chondroitin sulphate proteoglycans of PNNs, which has strong consequences on neuronal plasticity (Yang et al., 2021). In contrast with neuroestrogen functional effects limiting CA1 synaptic inhibition (Hernández-Vivanco et al., 2022), estrous cycle is not reflected in apparent changes in IPSCs frequency in CA1 pyramidal neurons or PV IN population activity monitored through fiber photometry, suggesting that brain-derived estrogen and ovarian hormones effect diverge in their functional effects in CA1 synaptic inhibition and PV INs.

The contribution and functional consequences of local synthesis of estrogen by hippocampal INs before sexual maturity, i.e. puberty, is currently unknown. Our results suggest that aromatase is expressed in CA1 PV INs in the male and female mice hippocampus at post-natal day 21. Thus, CA1 PV INs may contribute to local synthesis of estrogen before the start of sex hormone production by adult gonads. Interestingly, our results also show that systemic pharmacological blockade of aromatase activity in prepubertal mice has functional effects on synaptic inhibition of CA1 pyramidal neurons. Aromatase inhibition increases sIPSC frequency in CA1 pyramidal neurons of prepubertal female but not male mice. Since juvenile gonadal hormone production remains at very low levels, these results strongly suggest female-specific effects of brain-produced neuroestrogen in the physiology of hippocampal INs in prepubertal mice. Importantly, this sex effect is detected before functional maturation of gonads, again suggesting that neuroestrogen and ovarian hormones independently regulate the function of CA1 INs. Through the regulation of inhibitory signaling and PV INs PNNs in the prepubertal hippocampus, neuroestrogen may promote the refinement of network activity (Cossart and Khazipov, 2022), control the closure of inhibition-dependent critical periods for brain plasticity (Miranda et al., 2022) and promote in this way the formation of precise memories (Ramsaran et al., 2023). Moreover, neuroestrogen actions in the prepubertal brain may have a functional impact in the development and maturation of hippocampal INs that takes place during this stage of life, affecting processes such as programmed cell death and synaptogenesis (Lim et al., 2018; Wong and Marín, 2019).

Our results evidence functional effects of sex hormones in the neonatal mice brain. During this period, sex hormones and their receptors are at their highest levels, prior to gradually decline to adult levels (Turano et al., 2019). Early exposure to testosterone in female mice pups alters prepubertal hippocampal and PV IN function and renders adult CA1 inhibition and PV INs PNNs insensitive to neuroestrogen regulation. This suggests that neonatal production of testosterone by testes in male mice impacts neuronal activity in the early hippocampus and triggers organizational effects on CA1 synaptic inhibition and PV INs. Although not directly tested in the experiments presented here, aromatase expression in neonatal PV INs could support local aromatization to estrogen (Wu et al., 2009). Strikingly, no large sex differences have been detected in 17β-estradiol and testosterone concentrations in the neonatal hippocampus (Konkle and McCarthy, 2011). However, neonatal testosterone surge in male mice may alter the availability aromatase substrate (testosterone) and product (17β-estradiol) in a cell-type specific manner and cause sex differences by triggering transient or permanent effects in defined neuronal populations. Organizational effects of neonatal testosterone have been reported in hippocampal excitatory neurons and have been proposed to explain sex differences in estrogenic signaling through metabotropic glutamate receptor in hippocampal neurons in vitro (Meitzen et al., 2012). Thus, neonatal hormones may coordinately organize excitatory and inhibitory hippocampal neurons to promote sex specific regulation of excitatory / inhibitory balance in developing networks. Importantly, the actions of sex hormones on hippocampal inhibition described here coincide temporally with a critical period for Neurodevelopmental disorder (NDD) pathogenesis. Since INs function is compromised in NDD, the current findings suggest that gonadal hormones may regulate the impact of NDD related pathological alterations in INs.

The early life period coincides with the maturation and functional integration of different hippocampal neuronal types and the emergence and refinement of spatially-tuned activity characteristic of CA1 place cells (Wills et al., 2010) and hippocampal network activity synchrony (Farooq and Dragoi, 2019). The critical role of hippocampal INs in controlling spatial coding (Valero et al., 2022) and oscillations (Klausberger and Somogyi, 2008) raises the possibility of functional consequences of organizational actions of sex hormones in hippocampal processes known to be important for episodic memory. Moreover, by impacting neuronal communication, organizational actions of neonatal hormones may support functional network maturation and prevent deviations from normal neurodevelopmental trajectories with enduring deleterious consequences.

## Methods

### Animals

All experiments were performed according to protocols approved by the Institutional Animal Care and Use Committee of the Cajal Institute and by local veterinary authorities (Comunidad de Madrid). Group housed CD1 male and female mice were used for all experiments except for fiber photometry recordings, which were performed on C57BL/6J PV-Cre mice (*Pvalbtm1(cre)Arbr/J*). Mice were maintained in a 12 h light/dark cycle, 20–22 °C, 45–65% humidity and with unlimited access to food and water. All animals were obtained from the animal facility of the Cajal Institute. Age and sex of the animals is described for each experiment in the corresponding figure and legend.

### Estrous cycle monitoring

Estrous cycle was monitored by vaginal cytologies performed between 7 and 10 am. A vaginal lavage with 75 µl saline solution was collected using a P200 pipette with a rounded tip. The lavage was repeated several times to ensure efficient cell sampling and placed in a gelatin-coated microscope slide. After drying, the sample was stained with cresyl violet (0.1 %) and imaged in an optical microscope using 10x and 40x objectives. The estrous cycle stage was determined according to the relative presence of epithelial cells (nucleated and cornified) and leukocytes.

### Reagents and hormonal treatments

Letrozole (Tocris) was dissolved in DMSO to 12.5 mg/ml, further dissolved in saline solution to 62.5 μg/ml and administered at a dose of 0.5 mg/kg in intraperitoneal (i.p.) injections of 8 ml/kg. Testosterone propionate (Sigma) was dissolved in sesame oil by overnight magnetic stirring at a concentration of 2 mg/ml. Female pups received interscapular subcutaneous injections (50 μl) using a 25 gauge needle.

### Slice electrophysiology

To prepare acute slices for electrophysiological recordings, brains were quickly removed and coronal slices (300 µm) containing the dorsal hippocampus were obtained with a vibratome (4°C) in a solution containing: 234 mM sucrose, 11 mM glucose, 26 mM NaHCO3, 2.5 mM KCl, 1.25 mM NaH2PO4, 10 mM MgSO4, and mM 0.5 CaCl2 (equilibrated with 95% O2–5% CO2). Recordings were obtained at 30-32°C from CA1 stratum pyramidale neurons visually identified using infrared video microscopy in oxygenated artificial cerebrospinal fluid containing 126 mM NaCl, 26 mM NaHCO3, 2.5 mM KCl, 1.25 mM NaH2PO4, 2 mM MgSO4, 2 mM CaCl2 and 10 Mm glucose (pH 7.4). Patch-clamp electrodes contained intracellular solution composed of: 127 mM Cesium methanesulfonate, 2 mM CsCl, 10 mM HEPES, 5 mM EGTA, 4 mM MgATP, and 4 mM QX-314 bromide, pH 7.3 adjusted with CsOH (290 mOsm). GABA receptors mediated inhibitory spontaneous (sIPSCs) were registered by clamping neurons at 0 mV. Signals were amplified using a Multiclamp 700B patch-clamp amplifier and digitized using a Digidata 1550B (Axon Instruments, USA), sampled at 20 kHz, filtered at 10 kHz, and stored on a PC using Clampex 10.7 (Axon Instrumenst). Series resistance was monitored by a voltage pulse in every recorded cell and compared between experimental groups to discard effects due to recording conditions. IPSC were analyzed using pClamp (Axon Instruments) and a custom written software (Detector, courtesy J. R. Huguenard, Stanford University), as previously described (Manseau et al., 2010). Briefly, individual events were detected with a threshold-triggered process from a differentiated copy of the real trace. For each cell, the detection criteria (threshold and duration of trigger for detection) were adjusted to ignore slow membrane fluctuations and electric noise while allowing maximal discrimination of sIPSCs. Detection frames were regularly inspected visually to ensure that the detector was working properly.

### Fiber photometry recordings

Adult female PV-Cre mice were stereotaxically injected with adeno-associated viruses (pAAV.Syn.Flex.GCaMP6m.WPRE.SV40, serotype 1, Addgene) and custom-made optical fiber implants (0.39 Numerical Aperture, 400 µm core diameter, Thor Labs) were positioned above dorsal CA1 and firmly attached to the skull, as described previously (Hernández-Vivanco et al., 2022). Fiber position and AAV infection was verified histologically at the end of the experiment. Mice were habituated to the recording arena for 10 min (35 × 24 cm plastic enclosure in a soundproof container with constant illumination, 75 lux) for 5 days before recordings. On the recording days (3-4 weeks after surgery), mice were connected to a Tucker-Davis Technologies fiber photometry system and placed for 10 min in the enclosure. Mouse behavior was video-recorded and position tracked using DeepLabCut (Nath et al., 2019). Behavior-Depot software (Gabriel et al., 2022) was used to calculate instantaneous speed. Fiber photometry signals were processed using custom-made code with MATLAB (MathWorks) that can be found <u>here</u>. Signals were downsampled to 15Hz in accordance to behavioral frequency sampling video-recordings. After detrending and subtracting isosbestic signal (dF/F), robust z-score was calculated based on the median and the median absolute deviation for the complete recording. Alignment of the signals of interest for locomotion events, namely calcium dependent fluorescence and velocity, were performed using a custom-made MATLAB script. Threshold was set at 1cm/s to identify locomotion events. Finally, the speed modulation index (Fig. 1C) was calculated in each recording using a logarithmic fit of the curves defined as the mean of dF/F Z-score values as a function of binned speed.

### Contextual Fear Conditioning

Behavioral tests were performed during the light phase (7am -7pm) before weaning. Mice were habituated and handled by the experimenter (5 min / day, 3 days before training). During the training session on PND20, mice were allowed to freely explore the contextual fear conditioning cage (25×25 cm methacrylate cage with a metallic grid floor and scented with a 0.5 % ammonia) for 3 min. On min 4, three mild electric shocks (0.5 mA) lasting 2 seconds each were delivered through the metallic grid floor with 30 s inter-shocks intervals. After 1 additional min, mice were placed back into their home cages. Recall session was performed 1 day after training in the conditioning cage (no shocks, 5 min duration). During training and recall, mice behaviour was continuously recorded with a digital camera. Mice position, immobility and freezing were automatically determined with Any-maze software (Stoelting). Active fear responses (jumps and climbs) were visually determined by an experimenter blind to the condition tested.

### Tissue processing and immunohistochemistry

Mice were injected with a lethal dose of pentobarbital (150mg/kg) and perfused transcardiacally with cold PBS and 4% paraformaldehyde solution. Brains were extracted and submerged in fixative for 4 hours at 4º C. Coronal 40 µm thick vibratome sections containing dorsal hippocampus were blocked in PBS 0.3% BSA, 5% normal goat serum (NGS) and 0.3 %Triton X-100 followed by overnight incubation in PBS, 5% NGS and 0.3 %Triton X-100 with primary antibody: parvalbumin (guinea pig polyclonal, code GP42, Swant), cFos (rabbit polyclonal, code 226008, Synaptic Systems) and aromatase (in-house production, previously described and validated (Yague et al., 2006). Biotinylated Wisteria Floribunda Lectin (Vector Laboratories) was incubated in the same conditions as primary antibodies. After 3×15 minutes wash in PBST at room temperature, slices were incubated with 1:500 Alexa-conjugated secondary antibodies and Steptavidin (Alexa-Fluor 488, 555, Abcam) to reveal primary antibodies and biotynilated WFA, respectively. After 3 more step of washing in PBST, slices were mounted and covered on microscope slides using DAPI containing mounting medium.

### Image analysis

Images were obtained with a Leica SP5 confocal microscope (LEICA LAS AF software) using 20x or 40x objectives and 405, 488, 561 nm laser excitation wavelengths. 1024×1024 images with a resolution of 1.3-2.6 pixel/μm, at 2-4 µm step size were collected. Manually depicted ROIs delimiting CA1 PV neurons were used to determine fluorescence intensity in other channels (aromatase). For quantification of WFA staining, a lineal ROI surrounding PV neuron (3,8 µm width) was used. Mean pixel intensity in closed and lineal ROIs was determined in equally thresholded images. For c-Fos cuantification, background fluorescence was measured from manually selected location in acellular regions of the Stratum Oriens. In order to determine the number of c-Fos+ PV INs, the background corresponding to c-Fos images was multiplied by 1.5, 1.75 and 2.0 times (dynamic threshold) and subtracted from the corresponding c-Fos value in each individual PV IN. The fraction of cells above the dynamic threshold was determined in each individual mouse.

### Statistical analysis

All values are given in mean ±SEM, except when noted. Standard t tests were performed to compare Gaussian distributions while Mann-Whitney tests were used for non-gaussian distributions. One-or two-way ANOVA followed by Bonferroni’s post hoc test were used when noted.

For all tests, we adopted an alpha level of 0.05 to assess statistical significance. Statistical analysis was performed using Prism (Graphpad software).

## Notes

**Conflict of Interest** Authors report no conflict of interest

**Funding sources** This work was supported by a grant PID112428GB-I00 by MCIN/ AEI/10.13039/501100011033 to PM. AM-M is supported by a JAEIntro scholarship funded by CSIC. R V-G is supported by the Ph.D. fellowship PRE2021-099806 funded by MCIN/AEI/10.13039/501100011033 by “ESF Investing in your future”.

### Competing Interest Statement

The authors have declared no competing interest.

### Summary of Updates

Results section: Figure 1 was revised, new experiments added. Fig. 5 was added. The rest of manuscript sections have been modified accordingly.

